# Resistance to change? The impact of group medication on AMR gene dynamics during commercial pig production

**DOI:** 10.1101/659771

**Authors:** Jolinda Pollock, Adrian Muwonge, Michael R. Hutchings, Geoffrey Mainda, Barend M. Bronsvoort, Laura C. Duggan, David L. Gally, Alexander Corbishley

## Abstract

The anthropogenic selection of antimicrobial resistance (AMR) genes is under intense scrutiny, particularly in livestock production, where group antimicrobial administration is used to control disease. Whilst large epidemiological studies provide important data on the diversity and distribution of AMR genes, we have little insight into how group antimicrobial administration impacts AMR gene abundance and diversity within a system. Here, faecal microbiome and AMR gene dynamics were tracked for six months through a standard production cycle on a commercial pig unit. Our results demonstrate that specific AMR genes have reached an equilibrium across this farming system to the extent that the levels measured were maintained from suckling through to slaughter, despite increases in microbiome diversity in early development. These levels were not influenced by antibiotic use, either during the production cycle or following whole herd medication. Some AMR genes were found at levels higher than that of the bacterial 16S rRNA gene, indicating widespread distribution across the most common bacterial genera. The targeted AMR genes were also detected in nearby soil samples, several with decreasing abundance with increasing distance from the unit, demonstrating that the farm acts as a point source of AMR gene ‘pollution’. Metagenomic sequencing of a subset of samples identified 144 AMR genes, with higher gene diversity in the piglet samples compared to the sow samples. Critically, despite overwhelming and stable levels of resistance alleles against the main antimicrobials used on this farm, these compounds continue to control the bacterial pathogens responsible for production losses and compromised welfare.

**Importance:** Group antibiotic treatment has been used for decades to control bacterial diseases that reduce the productivity and compromise the welfare of livestock. Recent increases in antibiotic resistant infections in humans has resulted in concerns that antibiotic use in livestock may contribute to the development of untreatable bacterial infections in humans. There is however little understanding as to how the genes that bacteria require to become resistant to antibiotics respond during and after group antibiotic treatment of livestock, particularly in systems where high levels of antibiotics have been used for a prolonged period of time. We show that in such a system, levels of specific antibiotic resistance genes are high irrespective of group antibiotic treatments, whilst dramatic reductions in antibiotic use also fail to reduce the levels of these genes. These findings have important implications for public policy relating to the use of antibiotic in livestock farming.

## Introduction

Antimicrobials, including antibiotics and micronutrients such as copper and zinc (1), are used regularly in many agricultural systems worldwide to improve the health, welfare and productivity of livestock (2, 3). This has led to concerns about the anthropogenic selection of antimicrobial-resistant bacteria in livestock systems (3–5), particularly due to the potential transfer of antimicrobial resistance (AMR) genes from livestock to humans (6–8) and into the environment (9, 10).

In 2016, coordinated international action by the World Health Assembly, G7, G20 and the United Nations was called for following a wide-ranging review commissioned by the UK government, which included proposed global targets, based on livestock biomass, to reduce antibiotic use in livestock by 2025 (11). The European Commission currently benchmarks antibiotic usage between member states using a population correction unit (PCU), a standardised measure of livestock biomass (12). Whilst an imperfect measure, it is clear that the variability in antibiotic usage between countries (12, 13), agricultural sectors (14) and individual farms (15–17) is vast. Livestock antibiotic sales ranged from 2.9 to 453.4 mg/PCU across 30 European countries in 2016, with usage in the United Kingdom (UK) calculated as 45 mg/PCU (12). In Europe, tetracyclines, penicillins and sulphonamides accounted for 70% of livestock antibiotic sales (12), whilst in the UK, 52% of all antimicrobials sold for livestock were used in the pig and poultry sectors, with tetracyclines being reported as the highest sold antimicrobial class (14).

Given the substantial use of antimicrobials in pig and poultry production, the association between antimicrobial use and AMR has been an area of intensive study. Multiple studies have associated the use of antibiotics (13, 18–20) and zinc or copper supplementation (1, 21–24) in pig production with increases in phenotypic resistance in indicator organisms or pathogenic bacteria. Metagenomic data has revealed associations between antibiotic usage in pigs and poultry with increases in AMR gene prevalence, with lower AMR gene diversity, but higher AMR gene loads being observed on pig units in comparison to poultry units (25). Administration of oxytetracycline immediately post-weaning has also been shown to increase the abundance of AMR genes conferring tetracycline, beta-lactam and multi-drug resistance (26). The persistence of AMR genes on pig units has also been assessed, with differences in specific AMR gene levels being observed when comparing samples from sows and finishers (27).

There are currently few longitudinal studies of AMR in livestock systems, specifically for pig production (28). Those that exist for pig production are again predominantly focussed on specific indicator organisms (18, 20, 29–34), meaning that the resistance potential within the total microbiota is likely being underestimated (16, 32, 35). In addition, information on antimicrobial use is often derived from national figures (13, 25), rather than from individual units. There are currently no longitudinal studies describing the relationship between antimicrobial usage, microbiome diversity and AMR gene dynamics in pigs in a commercial setting. Such knowledge is key to defining the risk that antimicrobial use in pigs poses to contamination of the environment with AMR genes and in developing effective policies to minimise the proliferation and dissemination of AMR genes.

Here, a longitudinal study was carried out on a UK commercial pig unit with high antimicrobial usage to track variation in the faecal microbiome and AMR gene abundance and diversity. Piglets were sampled weekly from birth to slaughter over a six-month period that included the administration of acidified water, and three prolonged periods of in-feed antimicrobial administration (zinc, chlortetracycline and tylosin). Dry sows that were not administered in-feed antimicrobials were sampled in parallel, whilst soil sampling was carried out around the farm perimeter at a single time point to assess the abundance of potential AMR gene ‘pollutants’ in the environment. In addition, a final set of samples were taken from sows prior to, during and after a partial depopulation event (i.e. a management practice which lowers disease burden within a herd) which involved antibiotic administration to every pig on the farm to assess if this practice impacted on AMR gene abundance. To address our aims, 16S rRNA gene metabarcoding, quantitative PCR (qPCR) and whole genome shotgun metagenomics were used to measure microbiome diversity, AMR gene abundances and AMR gene diversity, respectively.

## Materials and methods

### Farm description

This study received ethical approval from the Royal (Dick) School of Veterinary Studies Veterinary Ethics Research Committee. A 600 sow Landrace x Large White commercial farrowing to finishing unit in the UK, with detailed antimicrobial usage records, was recruited. Batch farrowing occurred every four weeks, with the study starting one week prior to the October 2016 batch farrowing. Nursing sows and piglets were housed on slats, whilst dry sows were housed in straw yards. Additional herd information is available in **Supplementary Materials 1**.

### Sampling

To capture gut microbiota and AMR gene dynamics on this farm, we collected weekly faecal samples (no sampling on week 6 (W6) due to access issues) starting on 26^th^ October 2016 and ending on 5^th^ April 2017 when the studied batch of pigs were sent for slaughter. The study design, sampling points and antimicrobial agents used are visualised in **Fig 1**. On W1, faecal drop samples were collected from the floor of six farrowing crates containing pregnant sows (n = 6). Between W2 and W4, both sow and piglet faecal drop samples were obtained from the same farrowing crates as W1. On W5, all piglets were weaned and mixed into three groups prior to movement into the weaning house. From W5 to W13, pooled faecal drop samples were taken from each of these three pens to capture pen-level dynamics. On W14, these pigs were moved into the grower/finisher house and remained in the same pen formation as in the weaner house. Thereafter, from W14 to W25, pooled faecal drop samples were taken weekly from each of these pens until slaughter.

**Figure 1.**
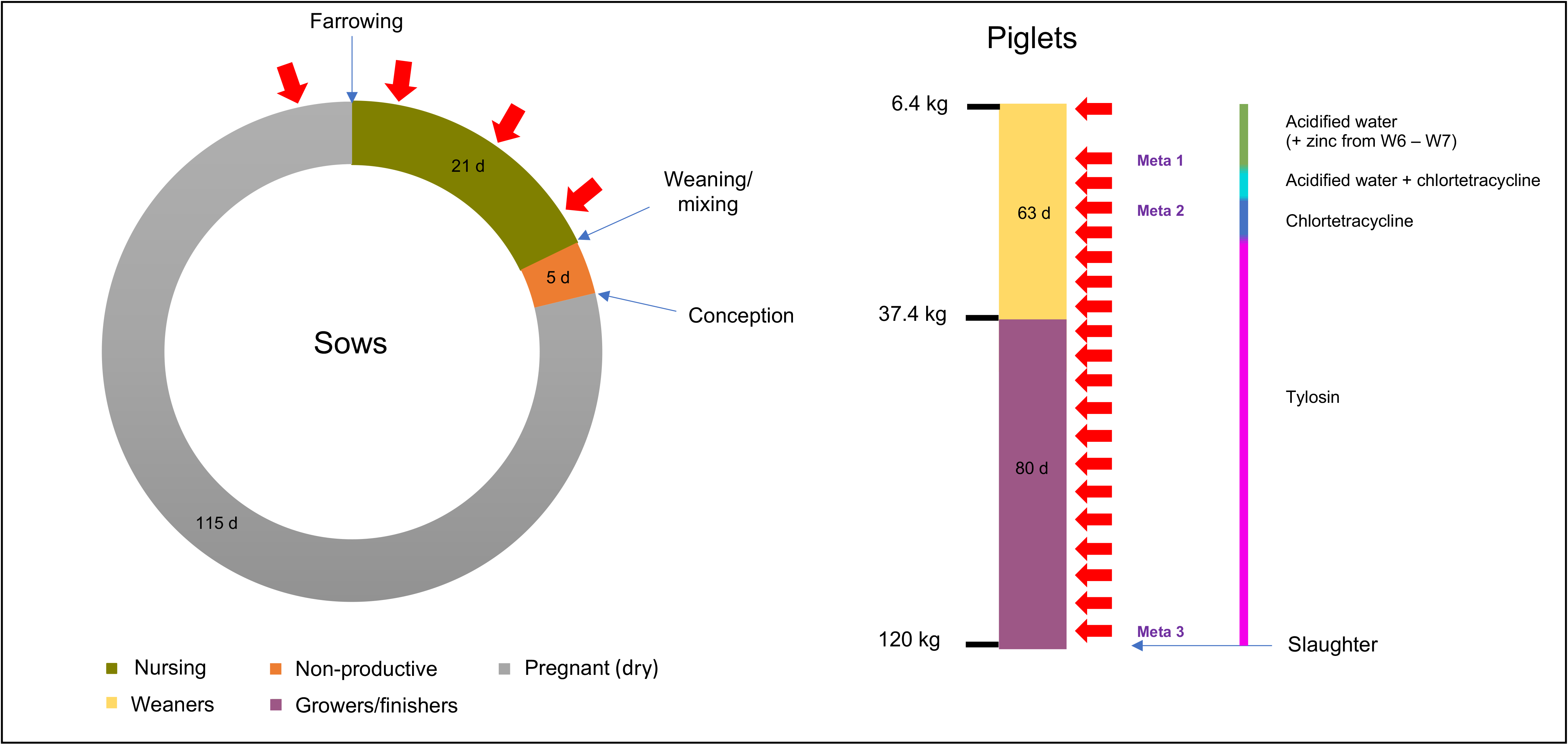
Summary of pig production cycle, antimicrobial administration and sampling points. Faecal sampling (red arrows) was carried out from farrowing pens containing the pregnant sows only (W1), and then post-farrowing from both the sow and the piglets during co-housing (W2-W4). Weaners were mixed and moved into new accommodation prior to sampling on W5 and were moved to larger accommodation for the growing/finishing period prior to sampling on W14. Samples subject to metagenomic sequencing are indicated (Meta 1 to Meta 3).

Weekly faecal samples were also taken randomly from the sow barn (n = 6) and slurry tanks (n = 2) to establish background levels of resistance in a group of adult pigs not given in-feed antimicrobials and across the farm as a whole, respectively. To examine the impact on the immediate surroundings of this farm, we also took samples from the environment around the pig unit. Thirty sampling locations were randomly chosen from within the farm boundary using the geographic information system application QGIS (QGIS Development Team, Version 2.18.13). Samples were taken from the surface at each point, and the substrate type was recorded (i.e. soil or faeces). If a sampling point was unsuitable due to features on the ground (i.e. standing water, access issues or trees), the point was moved the minimum distance necessary to clear the feature in a random direction and the new co-ordinates were recorded. As the study farm was located in a region of high pig farm density, comparator soil samples were obtained from areas close to the investigators’ laboratory that had no history of exposure to agricultural slurry (i.e. public parks and forestry areas).

In an effort to improve health status and reduce antimicrobial use, the farm underwent a partial depopulation in May 2017. This involved removing all the young pigs (i.e. sucking piglets, weaners, growers and finishers) from the farm and fumigating the young pig accommodation with formaldehyde. Faecal samples were obtained from the sow barn during (W36) and after (W45 and W57) (n = 6) antimicrobial treatment of the sows used as part of the partial depopulation.

All samples described were collected in spooned universal tubes and stored at −20°C on site and samples were batch transported back to the laboratory on dry ice before being stored at −80°C prior to processing.

### DNA extractions and dry matter calculations

All DNA extractions were carried out using the DNeasy PowerSoil Kit (Qiagen, UK). Briefly, 250 mg of faeces or soil were thoroughly homogenised and then weighed prior to being transferred into the included bead tube with 60 μl of Solution 1. The samples were then mixed using a vortex prior to being homogenised for 45 seconds at 5.0 m s^-1^ using a FastPrep FP120 Cell Disrupter (Qbiogene Inc, France). The homogenised samples were then processed according to the manufacturer’s instructions.

In parallel, approximately 1 g of homogenised faeces or soil were weighed out, transferred to a foil weighing boat and placed into a drying oven set at 60°C overnight. In order to calculate the percentage of dry matter (% DM), the foil weighing boats were weighed prior to and after the material was added and were re-weighed after overnight drying.

### Quantitative PCR

Five AMR genes were selected to quantify using qPCR. The selection of these genes was based on the results of an initial end-point PCR screening of a sub-sample of DNA extracts obtained from the final sampling point (faecal samples = 6, soil samples = 3). A panel of 30 genes were selected, on the basis that these genes were of biological relevance to the historic use of antimicrobials on the pig unit and of importance to both veterinary and human medicine (see **Supplementary Materials 2** for list of target genes and primer sequences). Genes which were amplified from more than 50% of the faecal samples were shortlisted for qPCR analysis. The selection for qPCR contained genes associated with tetracycline (*tetB* and *tetQ*), tylosin (*ermA* and *ermB*) and trimethoprim resistance (*dfrA1*) which are linked to the antimicrobials historically and currently administered in-feed on the farm. Quantification of the 16S rRNA gene was included as a proxy of overall bacterial load. Plasmids containing gene fragments of *tetB*, *tetQ*, *ermA*, *ermB*, *dfrA1* and the 16S rRNA gene were generated in-house for absolute quantification of gene copy number.

qPCR mixtures were set up using Brilliant III Ultra-Fast qPCR Mastermix (Agilent Technologies, United States), reference dye (Agilent Technologies, United States) and the primers and probes (as designed previously (35–50) or in the current study) listed in **Supplementary Material 2**. Each reaction was carried out in triplicate in a final volume of 20 μl, containing 1 μl of extracted DNA. Twenty-four samples were run per 96-well plate, which also included DNA standards (at concentrations ranging from 10^7^ to 10^1^ gene copies per μl) and a no-template control (nuclease-free water). Absolute quantification was carried out using a Stratagene MX3005P qPCR System (Agilent Technologies, UK) using the following fast, two-step cycling conditions: 95°C (5 minutes), followed by 40 cycles of amplification at 95°C (15 seconds) then 60°C (30 seconds).

Standard curves were constructed from the threshold cycle (C_T_) values using the Stratagene MxPro Software (Agilent Technologies, UK) and within this software, the calculated gene copy number per μl for each of the samples was generated, treating each of the three replicates individually. Any samples which fell beneath the limit of detection (i.e. 10^1^ copies per µl DNA, equivalent to 3.3 – 4.9 log_10_ copies/g DM) were re-run to confirm the findings. These values were then exported in Microsoft Excel spreadsheet format and the arithmetic means for the technical replicates calculated. These values were then converted into gene copy number per gram of dry matter (using the % DM values calculated) and log_10_-transformed for data visualisation and statistical analysis.

### 16S rRNA gene metabarcoding

All samples collected from the sow barn and piglet accommodation were prepared for 16S rRNA gene metabarcoding targeting the V3 hypervariable region, as described in previous work (51). Six library pools were compiled using equimolar concentrations of DNA from each sample. A mock bacterial community (20 Strain Even Mix Genomic Material ATCC®MSA-1002, ATCC, United States) and a reagent-only control (generated by passage of nuclease-free water throughout the library preparation process) were included in each pool to assess background contamination and sequencing error rate. Using the mock bacterial community sequences, the mean sequencing error rate was calculated as 0.01%.

The pools were submitted to the sequencing centre (Edinburgh Genomics, United Kingdom) where the pools were quantified using the Quant-iT™ PicoGreen® double-stranded DNA Assay Kit (Thermo Fisher Scientific, UK) to ensure sufficient yield for sequencing. Sequencing was carried out using the Illumina MiSeq platform (Illumina, United States), using V2 chemistry and producing 250 bp paired-end reads. The sequence files generated, with the primers removed, will be made publicly available through the NCBI Sequence Read Archive (SRA).

The generated sequences were processed using cutadapt (Martin, 2011) and mothur (Schloss et al, 2009) (URL: https://www.mothur.org/wiki/MiSeq_SOP. Accessed January 2018) as described previously (51). Here, unique sequences were binned using a database-independent approach. A mean of 125,312 sequences per sample were retained post quality control and were subsampled to 10,000 sequences per sample for analysis. The Inverse Simpson index (ISI) and Shannon index (SI) were calculated for each sample to assess alpha diversity.

### Statistical analysis

Repeated measures analysis of variance was carried out (Genstat 16, VSN International, UK) to assess temporal changes in gene copy number or alpha diversity indices in both the sow barn and piglet accommodation samples, with least significant differences being used for multiple comparisons of means to assess the impact of antimicrobial treatments on these parameters in the piglet accommodation. For the sow barn analyses, the values calculated from samples collected between W1-W25 were included in the statistical models. For the piglet accommodation analyses, the values calculated from samples collected between W5-W25 were included in the statistical models, since W1 samples were obtained from pregnant sows only and samples obtained between W2-W4 were obtained when the piglets were still grouped by litter and were not yet assigned to their rearing pens. For the piglet accommodation analyses, the pen was included as a factor to assess any differences between the triplicate pen samples.

### Shotgun metagenomic sequencing

Three time points were selected that corresponded to antimicrobial treatment in the piglets (**Fig 1**). Triplicate samples from both the sow barn (i.e. no in feed antimicrobial use) and piglet accommodation were submitted for shotgun metagenomic sequencing (Edinburgh Genomics, United Kingdom). Illumina TruSeq DNA Nano libraries were prepared using the submitted faecal DNA extracts and sequencing was carried out using the HiSeq 4000 platform generating 150 bp paired-end reads (Illumina, United States).

Host DNA was removed from the raw reads by mapping to the *Sus Scrofa* reference genome GCA_000003025 version 11.1 (52) and Phix DNA (PhiX 174) (53) was removed using the run_contaminant_filter.pl script which is part of the Microbiome Helper suite (54). Read quality control was carried out using trimmomatic (55). These paired end reads were then mapped to MEGARes (56), a hand-curated AMR gene database, using the paired-end option of BWA (57). The resistome profiling was carried out using the ResistomeAnalyzer function in MEGARes with the gene fraction threshold set at 90. In order to obtain a read per kilobase (RPK) normalised abundance count, the Humann2 script which is part of the Humann2 pipeline (58) was used to analyse the SAM file produced from the mapping step above. The resultant gene family output was then normalised using the humann2_renom_table script. The generated tsv and csv files were then used to analyse AMR gene abundances in R version 3.5 (59). The downstream analysis of AMR gene diversity and data visualisation were carried out using the ggplot2 (60) package in R.

## Results

### Defining antibiotic usage

In the three months prior to the study period, a total of 389.1 mg/PCU of antibiotics were used on the study farm. This compares to a UK average in pigs of 183 mg/PCU in 2016 (61), making this a high antibiotic usage farm. During the study period, the following routine group medication regimens were used (**Fig 1**): toltrazuril (30 mg/head oral) at 4 days old to control *Isospora suis*, zinc (2500 ppm in feed) between 4 and 6 weeks old to control post-weaning colibacillosis, acidified water (Baynes Evacide S 0.2%) to control post-weaning colibacillosis between 3 and 7 weeks old, chlortetracycline (300 ppm in feed) from 6 to 8 1/2 weeks old to control *Mycoplasma hyopneumoniae* and *Mycoplasma hyorhinis* and tylosin (100 ppm in feed) from 8 1/2 weeks old until slaughter to control *Mycoplasma hyopneumoniae*, *Mycoplasma hyorhinis*, *Actinobacillus pleuopneumoniae* and *Lawsonia intracellularis*.

During the partial depopulation, the dry sows received 1500 ppm chlortetracycline and 500 ppm tiamulin in-feed, whilst the nursing sows received 1875 ppm chlortetracycline and 625 ppm tiamulin. In the three months that included this partial depopulation, total antimicrobial use increased to 582.8 mg/PCU, which then declined to 32.3 mg/PCU in the three months after the partial depopulation.

### AMR gene abundances and microbiome diversity

All five genes studied reached levels exceeding 7 log_10_ copies/g of dry matter (DM) faeces in both the piglet accommodation and sow barn during the study (**Fig 2**). Prior to weaning, the levels of all five genes were similar when comparing the nursing sow and piglet samples (**Fig 2a**), even though the alpha diversity of the faecal microbiome was markedly lower in samples obtained from the piglets (**Fig 3**). As the alpha diversity increased in the piglet samples over the first few weeks of life, the AMR gene counts remained relatively stable, despite marked restructuring of the faecal microbiota.

**Figure 2.**
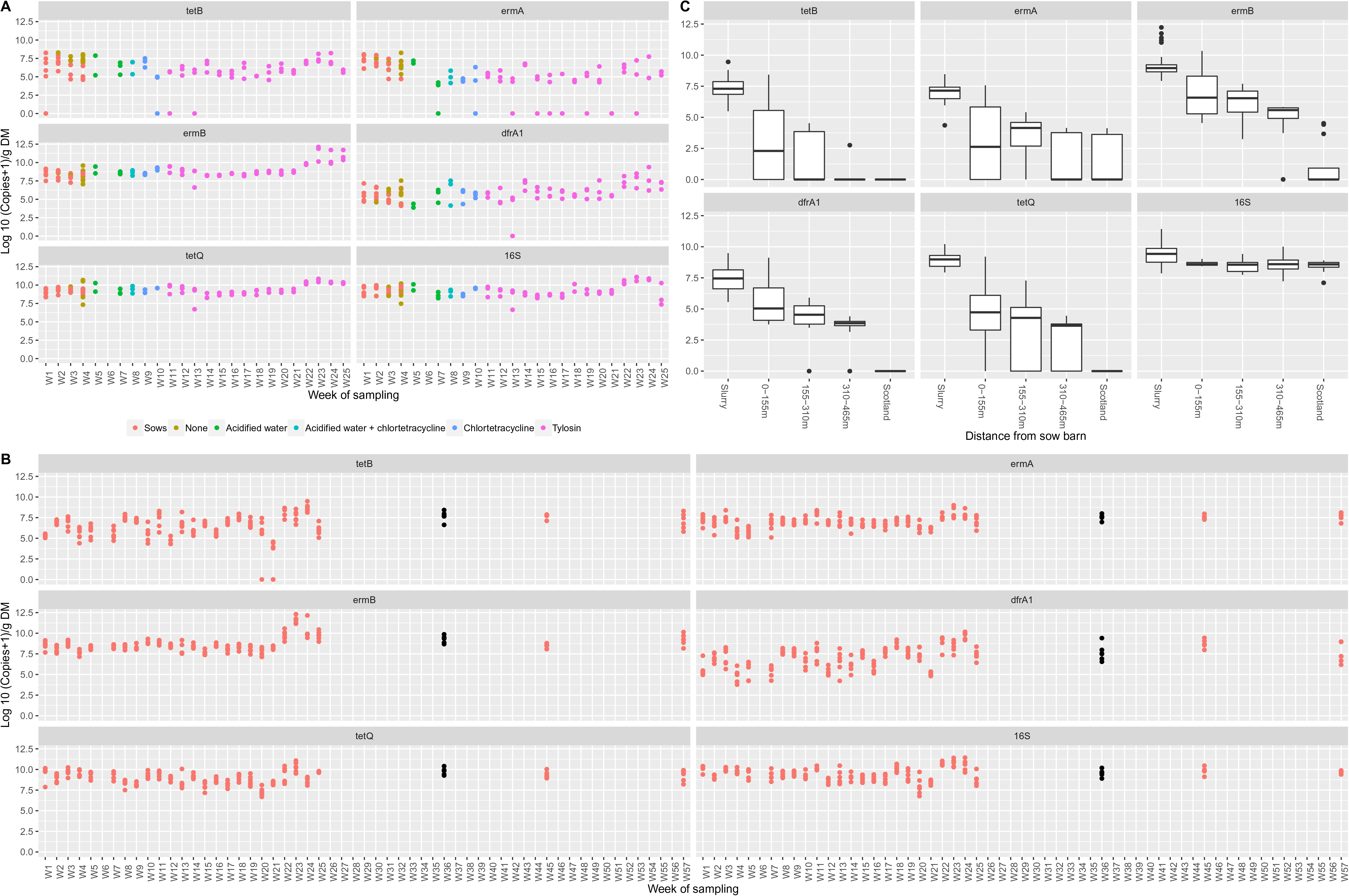
Quantification of AMR genes. AMR gene copy numbers in faecal samples obtained from the (A) piglet accommodation, (B) sow barn including the partial depopulation and (C) local environment, including slurry and soils obtained from non-agricultural sites in Scotland.

**Figure 3.**
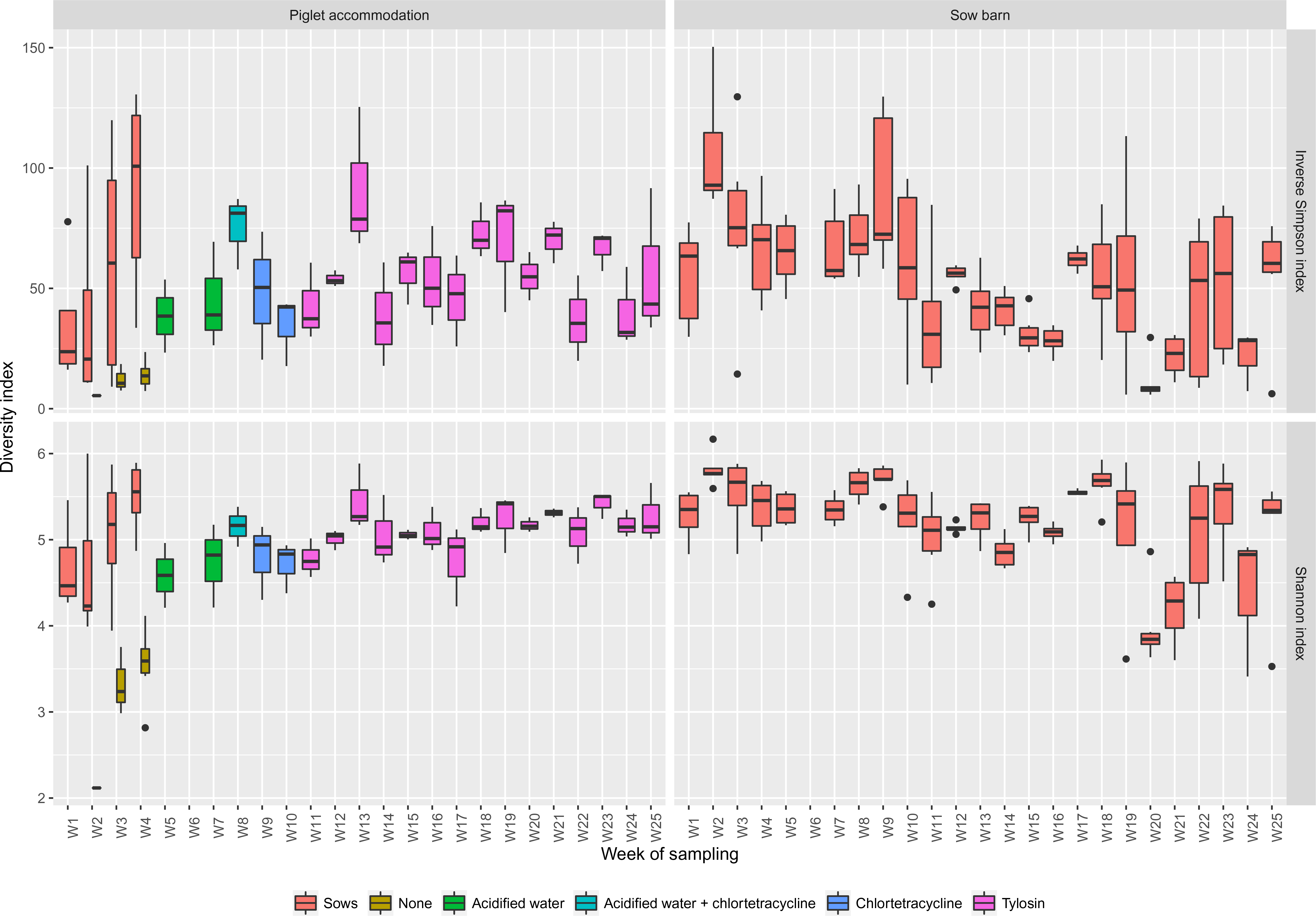
Alpha diversity indices of the faecal microbiota. The Inverse Simpson and Shannon indices of the faecal microbiomes obtained from the piglet accommodation and sow barn samples. On W1, faecal samples were obtained from pregnant sows only (n = 6) and on W2-4, faecal drops were obtained from both nursing sows (n = 6) and piglets (n = 6) during co-housing.

When comparing AMR gene abundances in the piglet accommodation samples (**Fig 2a**) after weaning (W5-W25) to the sow barn samples (**Fig 2c**), *tetQ* was the most abundant gene in both locations, whilst *ermA* and tet*B* were the least abundant in the piglet accommodation and sow barn, respectively. At times, the levels of *tetB*, *ermA* and *dfrA1* dropped below the limit of detection (3.3 – 4.9 log_10_ copies/g DM, depending on the DM of the sample) for particular samples. The 16S rRNA gene was detectable at levels exceeding 6 log_10_ copies/g DM faeces in all samples and was similar when comparing the piglet accommodation (mean 9.18 log_10_ copies/g DM, standard deviation (SD) 0.89) and sow barn samples (mean 9.41 log_10_ copies/g DM, SD 0.84), indicating that bacterial DNA was detectable in all samples processed and bacterial load was similar in piglet and sow faeces.

Despite the sows receiving no group antimicrobial treatments during W1-W25, levels of *tetB*, *ermA* and *dfrA1* were numerically higher in the sow barn (means 6.39, 6.93 and 6.98 log_10_ copies/g DM respectively, SD 1.50, 0.74, 1.44) compared to the post-weaning piglet accommodation (means 5.81, 4.24, 5.96 log_10_ copies/g DM respectively, SD 1.61, 2.33, 1.38). Conversely, levels of *tetQ* and *ermB* were numerically higher in the post-weaning piglet accommodation (means 9.18, 9.02 log_10_ copies/g DM respectively, SD 0.73, 1.01) compared to the sow barn (means 8.91, 8.62 log_10_ copies/g DM respectively, SD 0.80, 0.96).

All the AMR genes were subject to temporal fluctuations in both the piglet accommodation and sow barn samples, with significant changes in 16S rRNA gene copy number also occurring in both locations (**Table 1**). There were no significant differences in AMR and 16S rRNA gene levels (***P* > 0.05**), Shannon indices (SIs) (***P* = 0.86, *F* = 0.15**) or Inverse Simpson indices (ISIs) (***P* = 0.50, *F* = 0.72**) when comparing triplicate pen samples from the piglet accommodation between W5 and W25.

**Table 1.** Summary of statistical model outputs (F-statistics and P-values) for assessing temporal shifts in gene abundances and microbiome diversity. ISI = Inverse Simpson Index. SI = Shannon Index.

|  | Sow barn |  | Piglet accommodation |  |
| --- | --- | --- | --- | --- |
|  | F-statistic | P-value | F-statistic | P-value |
| *tetB* | 12.16 | < 0.001 | 2.05 | 0.030 |
| *tetQ* | 11.83 | < 0.001 | 5.60 | < 0.001 |
| *ermA* | 7.19 | < 0.001 | 2.01 | 0.034 |
| *ermB* | 30.02 | < 0.001 | 8.94 | < 0.001 |
| *dfrA1* | 20.69 | < 0.001 | 2.59 | 0.006 |
| **16S rRNA** | 13.30 | < 0.001 | 4.84 | < 0.001 |
| **ISI** | 5.63 | < 0.001 | 1.79 | 0.065 |
| **SI** | 5.79 | < 0.001 | 1.78 | 0.065 |

The greatest change in AMR gene prevalence occurred during W22-24 (**see Supplementary Materials 3** for statistical output), during which time the abundance of all five genes studied increased in both the piglet accommodation and sow barn (**Fig 2a and 2b**). This coincided with an increase in the bacterial 16S rRNA gene abundance, suggesting an increase in overall bacterial load across both locations and was independent of antimicrobial administration, given that the sows were not receiving any treatment and the piglets had been on tylosin continuously for 12 weeks prior to this point.

Other notable changes in AMR gene copy number include an increase in *dfrA1* counts in the piglet accommodation during acidified water and zinc administration between W5 (mean 4.17 log_10_ copies/g DM, standard error of the mean (SEM) 0.80) and W7 (mean 5.58 log_10_ copies/g DM, SEM 0.65) and a decrease in *ermA* between W5 (mean 6.83 log_10_ copies/g DM, SEM 1.44) and W7 (2.70 log_10_ copies/g DM, SEM 1.17). There were no marked shifts in the levels of *tetB, ermB* and *tetQ* between W5 and W7. A decrease in *tetB* gene copy number occurred in the piglet accommodation between W9 (mean 6.95 log_10_ copies/g DM, SEM 0.79) and W10 (mean 3.29 log_10_ copies/g DM, SEM 0.79) with no significant changes in *tetQ, ermA, ermB or dfrA1* levels being observed, despite the piglets having already been on chlortetracycline treatment for three weeks by W10.

Tylosin administration (W11-W25) did not have an effect on AMR and 16S rRNA gene counts in the piglet accommodation following withdrawal from chlortetracycline, but when the pigs were moved from the weaner/grower house (W13) into the finisher house (W14), increases in both *dfrA1* (3.37 to 6.99 log_10_ copies/g DM, SEM 0.65), *ermA* (3.03 to 6.69 log_10_ copies/g DM, SEM 1.17) and *tetB* (3.84 to 6.50 log_10_ copies/g DM, SEM 0.79) occurred, with *ermB* and *tetQ* counts not changing markedly over this period.

In the sow barn samples, similar to the AMR genes, both SIs and ISIs varied significantly over time, with the piglet accommodation samples between W5-W25 remaining more stable and did not vary significantly during this phase (**Table 1**). There were, however, notable stepwise increases in both SIs (from 2.12 (W1) to 4.59 (W4), SD 0, 0.53) and ISIs (from 5.43 (W1) to 38.47 (W4), SD 0, 21.46) in the piglet accommodation samples during the first 4 weeks of life. In both locations, both the SIs and ISIs appeared unstable and oscillated between W8 and W25, which emphasises changes in the number of unique sequences and large variations in the number of sequences assigned to each taxonomic group, respectively.

### AMR gene diversity

Triplicate samples from three time points were submitted from the sow barn and piglet accommodation for metagenomic sequencing. Across the 18 samples, a total of 144 AMR genes were identified (**Fig 4a**), 21 of which were ubiquitous across sampling location and time. The majority of the 60 genes that were present in both the piglet accommodation and sow barn were more abundant in the piglet accommodation. That said, abundance was generally within the same order of magnitude between the two sampling locations, whilst there were also notable examples where abundance was higher in the sow barn compared to the piglet accommodation. In agreement with the qPCR results, two of these genes were *dfrA1* and *ermA*. Also in agreement with the qPCR results, *tetQ* and *ermB* were more abundant in the piglet accommodation than the sow barn. In contrast to the qPCR results, *tetB* was more abundant in the piglet accommodation. There was noticeably more diversity in AMR genes within the piglet accommodation compared to the sow barn, with 77 genes found solely in the piglet accommodation, compared to just 7 genes that were unique to the sow barn.

**Figure 4.**
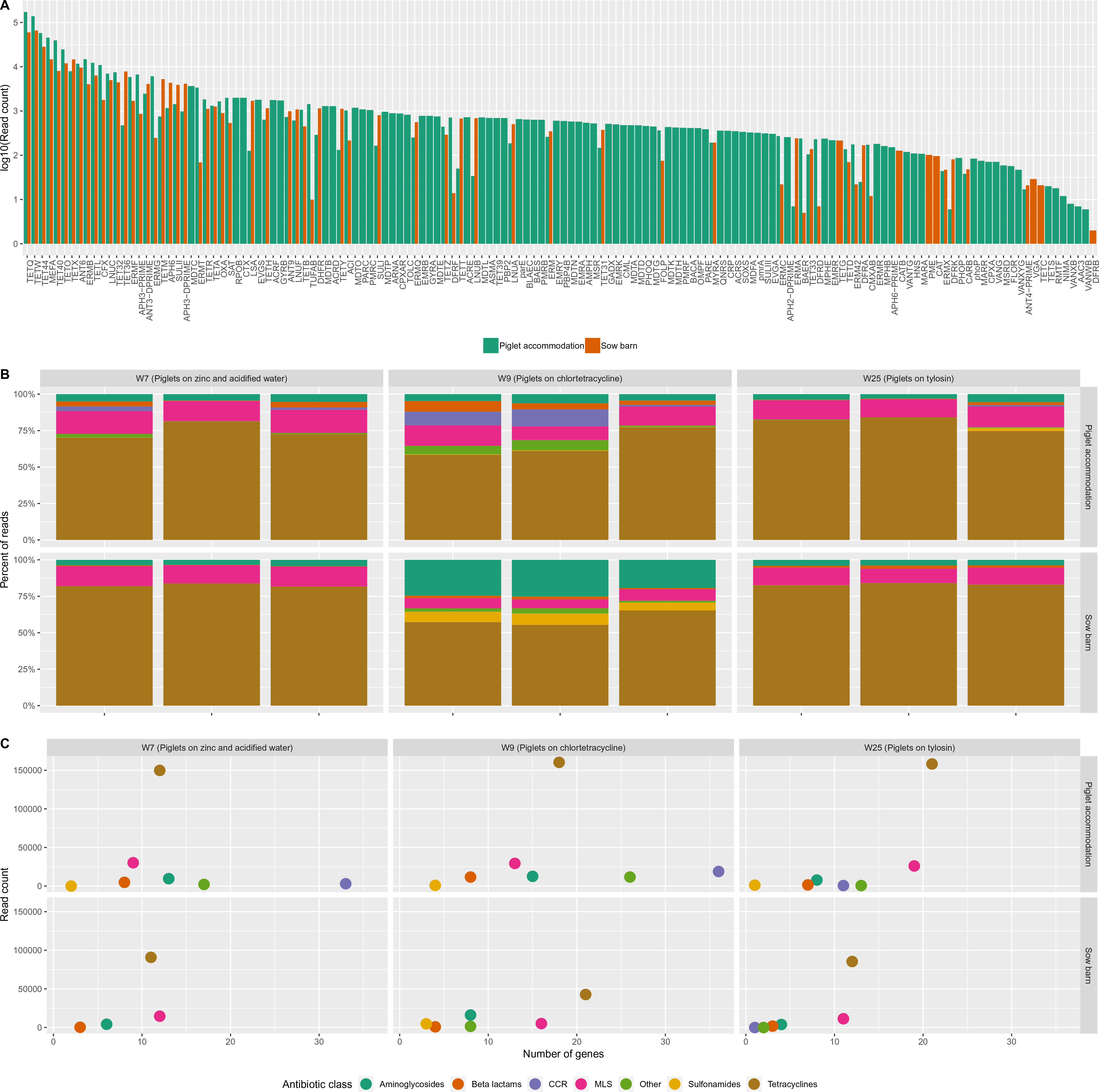
AMR gene abundance and diversity described by metagenomic data. (A) Log-transformed normalised gene abundance counts from both the sow barn and piglet accommodation (each bar is the sum of nine samples), (B) relative abundances of genes conferring resistance to particular antibiotic classes and (C) the sum of the absolute read counts from the three replicate samples conferring resistance to particular antibiotic classes.

When examining the proportion of reads associated with each antimicrobial class, AMR genes associated with tetracycline, macrolide (MLS) and aminoglycoside resistance predominated across all samples (**Fig 4b**). With a few exceptions, these proportional results were consistent across the triplicate biological replicates. These proportional results were also broadly similar for the sows and piglets on and between W7 and W25, despite the piglets having been on tylosin treatment for a number of months at the W25 sampling.

The biggest change in the proportion of reads associated with each antimicrobial class occurred on W9 in both the piglet accommodation and sow barn. In the piglet accommodation, there was a proportional increase in reads associated with cross-class resistance (CCR). In the sow barn, the increase was seen in reads associated with aminoglycoside and sulphonamide resistance. These proportional changes were accompanied by an increase in AMR gene diversity from 94 genes in the piglet accommodation on W7 to 120 genes on W9 and 32 genes in the sow barn on W7 to 60 genes on W9. The proportion of reads associated with tetracycline resistance declined between W7 and W9, despite the piglets starting chlortetracycline treatment during this time. No change in management, feed or use of medications were reported in the sow barn between W7 and W9 to explain this shift in the relative abundance and diversity of resistance genes. These results suggest changes in AMR gene abundance and diversity in response to an as yet unknown factor that affected the whole farm, rather than a specific response to antimicrobial administration.

In the piglet accommodation, there were both high levels and numbers (i.e. > 30) of genes conferring cross-class resistance (CCR) when the piglets were being administered zinc and acidified water (W7) and chlortetracycline (W9), but this number dropped by the end of the production cycle, despite the piglets having spent several months on in feed antimicrobials (**Fig 4c**). Genes associated with CCR were not detected in the sow barn on W7 and W9, with only a single gene detected on W25. Nearly half i.e. 35 of the 77 genes found solely in the piglet accommodation were associated with CCR and appeared to be related to management interventions, potentially acidified water and zinc administration, other than in-feed antibiotic administration.

### AMR gene quantification during and after a partial depopulation

To improve the health status of the farm, all the young pigs were removed (partial depopulation) and the sows treated with in-feed chlortetracycline and tiamulin. During this time period, all the pigs on the farm were treated with antimicrobials simultaneously. Samples were taken from the sow barn during the last week of treatment (W36) and two (W45) and five (W57) months after cessation of treatment. Bacterial load and AMR gene copy number in the sow barn samples did not change either during or after in-feed antimicrobial administration (**Fig 2b**).

### AMR gene quantification in environmental samples

In the environmental samples (**Fig 2c**), the 16S rRNA gene copy number was lower when compared to the slurry samples, however it was stable across all the soil samples, regardless of their distance from the farm or location. In the comparator soils, there were no detectable levels of *tetB*, *tetQ* or *dfrA1* and low levels of *ermA* and *ermB* genes. The abundance of all five AMR genes was lower in the environmental samples, compared to the slurry samples, whilst *tetB*, *ermB* and *dfrA1* showed a clear dilution effect, with AMR gene copy number declining with increasing distance from the sow barn.

## Discussion

When designing this study, our first hypothesis was that there would be a reservoir of AMR genes within the faecal microbiome, which would increase in both abundance and potentially diversity in response to in-feed antimicrobial administration. Furthermore, we anticipated that AMR gene abundance would be inversely related to faecal microbiome diversity. Whilst large cross-sectional studies have been instrumental in furthering our understanding of AMR gene abundance and diversity across different livestock systems (13, 25, 62), they have not provided the granularity of data and medicines usage history required to test these hypotheses. For this reason, we chose to undertake a prolonged period of intensive sampling on a single large commercial pig unit during three ‘real world’ group in-feed antimicrobial treatment regimens.

### Measured AMR genes are at saturation in both medicated and non-medicated pigs

Although previous work has demonstrated baseline high levels of specific AMR genes (63) and phenotypic resistance in *E.coli* isolates (33, 64) in unmedicated pigs, our initial expectation was that AMR gene abundances would markedly increase in response to antimicrobial administration as a result of selective pressure being exerted on specific gene subsets (65). In agreement with previous work, we showed high levels of all studied AMR genes in the non-medicated sow barn population, whilst prolonged exposure to three different in-feed antimicrobial regimens had no marked effect on AMR gene counts, potentially suggesting that they have reached saturation within the faecal bacterial populations (65). Curiously, the antimicrobials used (chlortetracycline and tylosin) were still clinically effective on this farm, suggesting that despite a high abundance of resistance genes within the faecal microbiome, that they were not present, or at least not active, within the organisms of clinical interest, i.e. *Mycoplasma hyopneumoniae, Mycoplasma hyorhinis, Actinobacillus pleuopneumoniae and Lawsonia intracellularis*.

### Changes in AMR gene abundances were not associated with antimicrobial exposure

Our statistical models did demonstrate fluctuations in AMR gene abundance over time in both piglets and sows, however changes were not associated with antimicrobial administration. In fact, the largest increases in AMR gene abundance were seen across all genes and in both classes of stock at the same time, hence suggesting as yet undefined environmental influences on AMR gene abundance. A previous cross-sectional study highlighted that only 10-42% of the variation in AMR gene levels could be explained by factors included in statistical models (including lifetime antimicrobial exposure), suggesting that AMR gene levels are strongly influenced by a variety of other elements (16). Whilst these could be related to feed changes, the fact that the sows and piglets were housed differently and fed different diets (66) would suggest that this effect is due to some other factors affecting the entire farm, such as housing and management (62, 67, 68), ambient temperature (68) and/or humidity, changes in water supply or the introduction of an infectious agent.

### Microbiome diversity was not affected by antimicrobial exposure

Similar to the studied AMR genes, temporal changes were evident in alpha diversity indices in both sow barn and piglet accommodation samples. These changes were clearly more pronounced in the growing pigs, as the alpha diversity of the faecal microbiome increased from nursing to finishing, which has been shown previously (69, 70). However, changes associated with antimicrobial treatment were not observed. Previous work has demonstrated that sub-therapeutic administration of chlortetracycline and tylosin had no impact on alpha diversity indices (69, 71). Whilst changes in relative abundances of specific taxa have been observed in response to tylosin administration, it was previously reported that these shifts were temporary, suggesting that the gut microbiota post-weaning seems to be resilient to perturbation by antimicrobial agents (69). Despite using markedly higher levels of chlortetracycline and tylosin (300 ppm and 100 ppm, respectively), our results also found that antimicrobial treatment did not impact on microbiome diversity in the growing piglets.

### AMR gene abundances are high in nursing piglets with low microbiome diversity

With respect to our second hypothesis, even when the piglet faecal microbiome was at its least diverse during the suckling period, the AMR gene levels were comparable to that of the nursing sows. Although microbiome diversity increased dramatically in the piglets during the first few weeks of life, as reflected in previous studies (69, 72–75), this was not associated with changes in AMR gene prevalence. We expected, as others have proposed, that changes in the microbiota would influence AMR gene levels (27, 66, 76), but in fact the high levels of studied AMR genes in the young piglets reflected that of the sows. The most obvious explanation for this is a combination of vertical and horizontal transmission of bacteria at or shortly after birth (31, 77). The presence of comparable levels of AMR genes in both sows and piglets and the associated large differences in microbiome diversity suggest that the AMR genes studied appear to either be widespread across multiple taxa or highly concentrated within dominant taxa present throughout all stages of microbiota development.

### Metagenomics revealed high AMR gene diversity and cross-class resistance genes in medicated pigs

The shotgun metagenomics revealed a diverse set of AMR genes in the presence and absence of antimicrobial treatment, which reflects findings in other recent work (26). Specifically, genes associated with tetracycline and macrolide resistance predominated. This is not surprising given the history of high levels of tetracyclines and macrolide use on this farm. Reassuringly, there was no evidence of co-selection between antimicrobial classes following chlortetracycline or tylosin administration. This is a significant finding and relevant to the principles of antimicrobial stewardship, where veterinary surgeons are actively discouraged from using fluoroquinolones and 3^rd^/4^th^ generation cephalosporins, so as to minimise the risk that resistance to these critically important antibiotics in human medicine is selected for in livestock.

What was evident, were the large numbers of genes associated with cross-class resistance (CCR) in the piglet accommodation samples taken on W7 and W9 and how these reduced by W25 and were almost absent from the unmedicated sows. The presence of these genes was clearly not associated with the administration of chlortetracycline and tylosin in this study. Although the current study design does not allow us to disentangle temporal effects versus the effect of different treatments, it is interesting to note that the CCR genes were already highly abundant at W7, following a period of in-feed zinc administration and the use of acidified water. Copper and zinc salts are commonly administered in-feed at supranutritional levels due to their antimicrobial properties, with increasing doses of zinc oxide being previously shown to increase the abundance of both tetracycline and sulphonamide resistance genes in weaned pigs (78, 79).

The metagenomic sequencing results for the sow barn on W9 were unexpected. All three biological replicates demonstrated an increase in the proportion of reads associated with sulphonamide and aminoglycoside resistance. The diversity of AMR genes also doubled at this time point compared to W7, despite no antimicrobial administration to this group of pigs. An increase in AMR gene diversity was also seen in the piglet accommodation at the same time point, however the increase was seen in beta-lactam resistance genes, CCR genes and genes associated with other antimicrobial classes. This is unlikely to be an artefact, given that this observation was seen across two different locations at the same time and five of the six biological replicates. As with the qPCR results discussed above, this shift appears to be a consequence of an undefined environmental factor affecting the whole farm, however in this instance, the two groups of pigs are responding with different increases in gene diversity.

### Persistently high abundances of AMR genes after partial depopulation

The partial depopulation at the end of the study involved considerable antibiotic use, with every sow on the farm receiving in-feed tiamulin and chlortetracycline. The treated sows were followed for five months after this treatment, during a period where antibiotic use on the farm declined nearly twenty-fold. Despite such dramatic changes in antibiotic usage, there were no marked changes in AMR gene abundances. This therefore begs the question as to how long, if ever, it would take to see reductions in AMR gene levels following the reduction or cessation of antimicrobial use on farms with a previous history of high-level use.

### AMR genes as environmental ‘pollutants’

We also found that the AMR gene levels in the environmental samples at the furthest points from the farm were still markedly higher when compared to urban and non-livestock environmental soils. Spreading pig slurry onto fields is a potential route of transferring AMR genes into the environment and from pigs to humans, as the AMR gene levels in soil are increased (80, 81). The surrounding fields, from which we obtained several samples, had been spread with slurry one week earlier from the unit, which most likely had a role in elevating the AMR gene counts at these locations. Previous studies have shown the persistence of resistant bacteria originating from slurry for 300 days (81) and even up to 18 months (82), with markedly higher levels in agricultural soils than comparator soils (81–84). Our findings, combined with that from previous research, suggest that the farm is acting as a reservoir of AMR genes.

### Policy implications

The results of this study imply that once AMR genes become established within the microbiome, modest changes in antimicrobial use are unlikely to result in a significant reduction in gene carriage. Furthermore, once established, the continued use of the antimicrobials against which these genes confer resistance, is unlikely to make the situation with respect to AMR gene dissemination any worse. The priority should therefore be to prevent resistance becoming established to antimicrobial classes of critical importance to human health i.e. 3^rd^/4^th^ generation cephalosporins and fluoroquinolones. More generally, attempting to refine existing antimicrobial treatment regimens offers limited scope for reducing AMR gene carriage. Efforts to combat antimicrobial resistance should therefore focus on removing the necessity for widespread antimicrobial administration to livestock, which includes pursuing high health status herds, improved management, optimised nutrition and enhanced immune function.

## Conclusion

AMR gene abundance and diversity on this unit were high in both medicated and unmedicated pigs, likely as a consequence of prolonged antimicrobial usage, with the farm acting as a point source of AMR genes. In this context, in-feed antibiotic (chlortetracycline and tylosin) administration did not affect AMR gene abundance or diversity. Acidified water and zinc supplementation were linked to an increase in AMR gene diversity, which importantly included CCR genes, which decayed quickly after their withdrawal. The implications of this work are that AMR genes that are already established within the microbiome are likely to decay only slowly (if at all) following antimicrobial withdrawal and so efforts should focus on avoiding the use of critically important antimicrobials, where high levels of resistance are not already established. This study did not identify which bacteria carried these AMR genes or indeed which genes were being expressed. Future work therefore needs to determine in what organisms these genes are not only present, but also active.

## Acknowledgements

The authors would like to thank the farm manager, staff on the study farm and their veterinary surgeons for their support and assistance throughout the sample and data collection phase of this project.

