## Supplementary 1 for "Resistance to change? The impact of group medication on AMR gene dynamics during commercial pig production"

**Supplementary Materials 1 - Additional herd information**

The herd was known to be positive for the following diseases: Porcine Reproductive and Respiratory Syndrome (PRRS), *Mycoplasma hyopneumoniae*, *Mycoplasma hyorhinis*, *Lawsonia intracellularis*, *Actinobacillus pleuopneumoniae*, *Streptococcus suis* and *Haemophilus parasuis*.

Vaccines were used to control the following diseases: PRRS, erysipelas, parvovirus, clostridial disease, *Mycoplasma hyopneumoniae*, Porcine Circovirus Type-2 (PCV2) and enterotoxigenic *Escherichia coli*.

The following quantities of antimicrobials were used during the three months prior to the study period starting: 265.8 mg/PCU tylosin, 103.2 mg/PCU chlortetracycline, 9.0 mg/PCU dihydrostreptomycin, 5.6 mg/PCU benzylpenicillin, 3.4 mg/PCU amoxicillin, 1.3 mg/PCU lincomycin, 0.4 mg/PCU marbofloxacin, 0.3 mg/PCU oxytetracycline and 0.1 mg/PCU enrofloxacin.

Previous batches of piglets prior to the study period received an additional in feed antimicrobial treatment of trimethoprim (100 ppm feed) and sulfadiazine (500 ppm feed) between 4 and 6 weeks old to control *Streptococcus suis* and *Haemophilus parasuis*.

During the study period, a mean of 12.9 piglets were born alive per sow, with mortality to weaning of 11.1%.
