## Supplementary 3 for "Resistance to change? The impact of group medication on AMR gene dynamics during commercial pig production"

**Supplementary Materials 3 – Statistical outputs**

**Piglet accommodation:** Assessment of temporal variation of mean gene copy number and mean diversity indices by analysis of variance, presented with the mean standard error of difference (SED) for each model. The models excluded samples from W1 (pens only contained pregnant sows at this time point) and samples from W2-W4 when the piglets were still grouped by litter and were not yet assigned to the rearing pens.

| Week | 16S rRNA | *dfrA1* | *ermA* | *ermB* | *tetB* | *tetQ* | ISI | SI |
| --- | --- | --- | --- | --- | --- | --- | --- | --- |
| 5 | 9.736 | 4.168 | 6.827 | 9.021 | 6.577 | 9.735 | 39.16 | 4.579 |
| 7 | 8.591 | 5.577 | 2.702 | 8.598 | 6.240 | 9.068 | 44.94 | 4.736 |
| 8 | 8.971 | 6.244 | 4.968 | 8.606 | 6.433 | 9.386 | 75.45 | 5.156 |
| 9 | 8.678 | 5.535 | 4.561 | 8.434 | 6.952 | 9.242 | 48.18 | 4.797 |
| 10 | 9.546 | 5.561 | 3.597 | 9.162 | 3.293 | 9.578 | 34.46 | 4.716 |
| 11 | 9.212 | 5.660 | 5.266 | 9.173 | 3.782 | 9.519 | 42.75 | 4.777 |
| 12 | 9.040 | 5.211 | 5.063 | 8.746 | 5.891 | 9.420 | 53.82 | 5.008 |
| 13 | 8.295 | 3.371 | 3.032 | 8.106 | 3.838 | 8.465 | 91.04 | 5.442 |
| 14 | 8.829 | 6.992 | 6.694 | 8.235 | 6.497 | 8.461 | 38.11 | 5.057 |
| 15 | 8.431 | 5.947 | 3.238 | 8.224 | 5.525 | 8.729 | 56.37 | 5.056 |
| 16 | 8.667 | 5.746 | 3.180 | 8.491 | 5.359 | 8.890 | 53.61 | 5.092 |
| 17 | 8.494 | 5.759 | 1.798 | 8.282 | 5.944 | 8.873 | 45.84 | 4.754 |
| 18 | 9.789 | 5.473 | 4.464 | 8.665 | 5.092 | 9.045 | 72.84 | 5.204 |
| 19 | 8.989 | 5.879 | 3.545 | 8.693 | 5.524 | 9.134 | 69.58 | 5.240 |
| 20 | 8.987 | 6.190 | 5.077 | 8.587 | 6.190 | 9.088 | 55.04 | 5.170 |
| 21 | 9.047 | 5.484 | 0.006 | 8.723 | 5.680 | 9.198 | 70.13 | 5.311 |
| 22 | 10.344 | 7.384 | 6.167 | 9.794 | 6.930 | 10.290 | 36.93 | 5.076 |
| 23 | 10.979 | 7.663 | 4.187 | 11.373 | 7.517 | 10.659 | 66.67 | 5.419 |
| 24 | 10.777 | 7.707 | 5.798 | 10.545 | 7.310 | 10.335 | 39.79 | 5.178 |
| 25 | 8.523 | 6.974 | 5.531 | 10.917 | 5.787 | 10.272 | 56.26 | 5.274 |
| Mean SED | 0.492 | 0.928 | 1.670 | 0.443 | 1.136 | 0.376 | 16.32 | 0.757 |

**Sow barn:** Assessment of temporal variation of mean gene copy number and mean diversity indices by analysis of variance, presented with the mean standard error of difference (SED) for each model.

| Week | 16S rRNA | *dfrA1* | *ermA* | *ermB* | *tetB* | *tetQ* | ISI | SI |
| --- | --- | --- | --- | --- | --- | --- | --- | --- |
| 1 | 10.090 | 5.558 | 7.190 | 8.618 | 5.329 | 9.574 | 55.4 | 5.278 |
| 2 | 9.102 | 6.986 | 6.646 | 8.020 | 6.984 | 8.987 | 106.2 | 5.851 |
| 3 | 10.000 | 7.088 | 7.455 | 8.860 | 6.899 | 9.691 | 75.9 | 5.541 |
| 4 | 9.723 | 4.868 | 5.929 | 7.635 | 5.515 | 9.459 | 66.5 | 5.387 |
| 5 | 9.552 | 5.546 | 5.819 | 8.240 | 5.855 | 9.345 | 64.9 | 5.362 |
| 7 | 9.282 | 5.359 | 6.682 | 8.363 | 5.598 | 9.191 | 67.1 | 5.352 |
| 8 | 9.361 | 7.760 | 6.985 | 8.365 | 7.506 | 8.323 | 72.0 | 5.644 |
| 9 | 9.489 | 7.736 | 6.827 | 8.273 | 7.112 | 8.151 | 90.2 | 5.692 |
| 10 | 9.219 | 6.522 | 7.143 | 8.884 | 5.523 | 9.267 | 60.2 | 5.221 |
| 11 | 10.123 | 7.634 | 7.907 | 8.849 | 7.331 | 9.436 | 36.5 | 5.022 |
| 12 | 8.657 | 5.368 | 6.952 | 8.477 | 4.624 | 8.870 | 55.8 | 5.134 |
| 13 | 9.065 | 6.562 | 7.057 | 8.009 | 6.784 | 8.433 | 41.9 | 5.236 |
| 14 | 9.083 | 6.084 | 6.712 | 8.435 | 6.008 | 9.134 | 41.0 | 4.857 |
| 15 | 8.910 | 7.520 | 6.645 | 7.632 | 6.794 | 7.925 | 31.4 | 5.248 |
| 16 | 8.845 | 5.811 | 6.615 | 8.487 | 5.603 | 8.640 | 28.3 | 5.084 |
| 17 | 8.760 | 7.771 | 6.606 | 8.067 | 7.025 | 8.209 | 61.6 | 5.550 |
| 18 | 10.109 | 8.570 | 7.152 | 8.327 | 7.593 | 9.085 | 54.2 | 5.650 |
| 19 | 9.213 | 7.616 | 7.086 | 8.007 | 6.756 | 8.634 | 54.5 | 5.085 |
| 20 | 7.987 | 7.506 | 6.352 | 7.728 | 5.248 | 7.273 | 11.4 | 3.980 |
| 21 | 8.468 | 5.072 | 6.155 | 8.314 | 2.783 | 8.340 | 21.9 | 4.187 |
| 22 | 10.704 | 8.960 | 7.572 | 9.757 | 8.211 | 9.278 | 44.7 | 5.073 |
| 23 | 10.791 | 8.420 | 8.044 | 11.611 | 7.607 | 10.263 | 52.9 | 5.382 |
| 24 | 10.572 | 9.741 | 7.779 | 10.176 | 8.663 | 8.600 | 21.8 | 4.384 |
| 25 | 8.736 | 7.404 | 7.039 | 9.725 | 6.007 | 9.708 | 55.2 | 5.094 |
| Mean SED | 0.283 | 0.408 | 0.301 | 0.234 | 0.517 | 0.279 | 13.10 | 0.304 |
